# Genomic prediction offers the most effective marker assisted breeding approach for ability to prevent arsenic accumulation in rice grains

**DOI:** 10.1101/452276

**Authors:** Julien Frouin, Axel Labeyrie, Arnaud Boisnard, Gian Attilio Sacchi, Nourollah Ahmadi

## Abstract

The high concentration of arsenic in the paddy fields and, consequently, in the rice grains is a critical issue in many rice-growing areas. Breeding arsenic tolerant rice varieties that prevent *As* uptake and its accumulation in the grains is a major mitigation options. However, the genetic control of the trait is complex, involving large number of gene of limited individual effect, and raises the question of the most efficient breeding method. Using data from three years of experiment in a naturally arsenic-reach field, we analysed the performances of the two major breeding methods: conventional, quantitative trait loci based, selection targeting loci involved in arsenic tolerance, and the emerging, genomic selection, predicting genetic values without prior hypotheses on causal relationships between markers and target traits. We showed that once calibrated in a reference population the accuracy of genomic prediction of arsenic content in the grains of the breeding population was rather high, ensuring genetic gains per time unite close to phenotypic selection. Conversely, selection targeting quantitative loci proved to be less robust as, though in agreement with the literature on the genetic bases of arsenic tolerance, few target loci identified in the reference population could be validated in the breeding population.

## Introduction

A survey of total arsenic (*As*) in 901 samples of commercial polished (white) rice collected randomly from arsenic contaminated or non-contaminated areas in 10 countries showed 7-fold variation in median total arsenic content. The lowest median value (0.04 mg/kg) was measured in Egypt and the highest in the U.S.A. and France, 0.25 and 0.28 mg/kg, respectively [1]. Pollution of paddy fields and irrigation water by *As* has been reported in more than 70 countries in Asia, America and Europe [2, 3]. The problem, which is often of geological origin, affects several hundred million peoples, especially in Asia [1, 3, 4]. Local and regional surveys revealed a tight correlation between *As* concentration in the soil, or in the irrigation water, and its concentration in the rice plant [2, 5]. At all sampling sites, *As* accumulation in the rice plant was the highest in the roots, followed by in the straw and cargo grain. Similar results have been observed in greenhouse experiments [6]. Pollution of the paddy field by *As* also affects crop growth and development (lower germination rate, reduced shoot and root growth and biomass production, etc.) and, consequently, crop yield [6].

Alternate wetting and drying of the paddy field during the cropping season is the most effective way of achieving agronomic mitigation [7]. Application of silicon (*Si*) fertilizer can also reduce the concentration of *As* in the rice plant [8]. A second category of mitigation options relies on rice genetic improvement to reduce *As* uptake and/or its translocation from the vegetative organs to the grains.

Mechanisms of rice plant response to soil *As* excess have been reported to be similar to those observed for other types of soil chemical toxicity [9]. However, the mechanisms related to the phytotoxic effects of *As* and the rice defense response to *As* remain poorly understood. In aerobic conditions, the predominant form of soil *As* is arsenate, *As*(OH)_5_ or *As*(V), and its uptake by plants involves phosphate transporters [10]. Overexposure to *As*(V) triggers reduced expression of genes coding for arsenate/phosphate transporters such as PHT1 [11]. At the same time, the arsenate taken up undergoes chemical reduction to a more highly toxic species, arsenite [*As* (III)] [12, 13]. The arsenite is then either excreted into the rhizosphere [14, 15], or transported to aboveground organs [16], and/or detoxified by complexation as phytochelatines and compartmentalized in the vacuoles [17]. In paddy fields, the predominant form of *As* is arsenite [18]. It enters root cells through aquaporin type membrane ports [19]. Transporters involved in the process include silicon transporters Lsi1 (influx) and Lsi2 (efflux) [19, 20] and several silicon-independent pathways [21, 22].

Significant genetic diversity for *As* accumulation has been reported in *A. thaliana* and in rice. In *A. thaliana*, genome-wide association analysis (GWAS) detected the HAC1 gene (High arsenic content 1) responsible for arsenate reductase activity in the root, facilitating arsenite efflux to the soil. In rice, significant genetic diversity for *As* accumulation has been reported under overexposure to *As* in both hydroponic cultivation and in field experiments [23–26]. Analysis of grain *As* content in 300 rice accessions grown in six sites distributed in Bangladesh, China and USA revealed from 3 to 34 fold variation in each site [25]. It also revealed that accessions belonging to the *Aus* genetic group had the highest *As* contents.

Using recombinant inbred lines (RIL) from bi-parental crosses, several QTLs involved in *As* accumulation have been mapped [23, 26–29]. Likewise, the use of phenotypic data produced in [27] for GWAS has detected several significant associations for grain *As* content [30]. However, none of the significant associations mapped in the vicinity (distance of less than 200 kb) of the Os02g51110 and Os03g01700 loci coding for Lsi1 and Lsi2 proteins, previously reported [19, 20] to play a central role in rice response to *As* overexposure. Likewise, very few significant associations colocalized with QTLs mapped in RIL populations [25]. Analysis of *As*-induced genome-wide modulation of transcriptomes of rice seedling roots revealed up-regulation of several hundred genes, confirming the complexity of the gene network involved in response to *As* overexposure [31–34]. Gene families with differential gene expression in *As* tolerant and *As*-susceptible genotypes include glutathione S-transferases, cytochrome P450s, heat shock proteins, metal-binding proteins, and a large number of transporters and transcriptions factors such as MYBs [35]. MYB genes may be crucial in *As*(V) stress tolerance as they upregulate phenylpropanoid and flavonoid biosynthetic pathways. More recently, using a reverse genetics approach, [36] showed that OsHAC1;1 and OsHAC1;2 (two orthologs of *A. thaliana* HAC1) functioned as *As*(V) reductases and played a role in the control of *As* accumulation in rice. Likewise, [14] showed that OsHAC4 played a critical role in rice tolerance to arsenate and regulated arsenic accumulation in rice. Based on these findings, some authors recently advocated using gene-editing technology to improve rice *As* tolerance [7, 22].

Here we report the results of our research into the potential of more conventional, marker assisted, breeding approaches to improve the ability of rice to restrict *As* accumulation in the grains. First, we used field phenotypic data (leaf and grain *As* content of rice plants grown on soil with rather high *As* concentration) and genotypic data from a reference diversity panel, to either map QTLs involved in *As* accumulation through GWAS or to train genomic prediction models. Second, using similar phenotypic and genotypic data from a panel of advanced lines from a breeding program, we analyzed congruence between GWAS results in the two populations, and evaluated the predictive ability of genomic prediction across the two populations. Our results identified genomic prediction as the most promising approach to improve the ability of rice to restrict *As* uptake and its accumulation in the grains.

## Results

### Phenotypic diversity for arsenic content

In 2014, soil analyses before crop establishment and after crop harvest revealed similar arsenic concentrations of about 10 mg kg^−1^ soil dry weight. During the same period, the monthly survey of the irrigation water revealed variable arsenic contents (0.014 to 0.034 mg l^−1^) with an average of 0.021 mg l^−1^. Similar soil and water arsenic contents were observed in 2015 and 2016 (S2 Table).

#### Variation in arsenic content in the reference population

The three arsenic-related traits evaluated exhibited normal distribution (Figure 1). Partitioning of the observed phenotypic variations into different sources of variation via the mixed model analysis revealed a highly significant effect of accession for all traits considered (Table 1). In 2014, the model R^2^ was greater than 0.70 for the three traits, indicating a good fit of the model. Similarly high R^2^ were observed in 2015 (0.63 for Ratio, 0.80 for FL-*As* and CG-*As*). Broad-sense heritability tended to confirm this trend, with values ranging from 0.80 to 0.86 in 2014, and above 0.91 in 2015 (Table 1).

**Figure 1:**
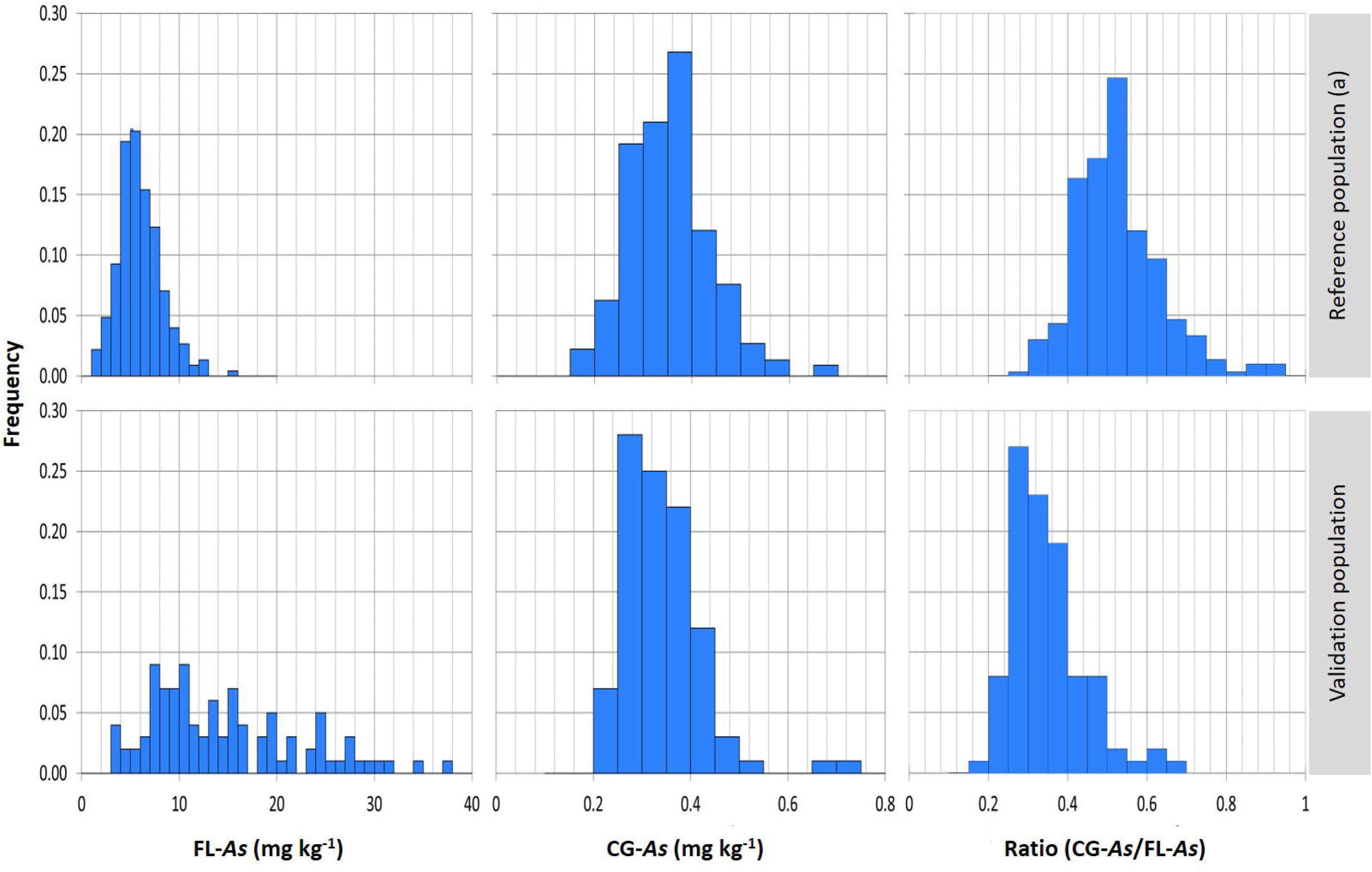
Distribution of adjusted phenotypic values for flag leaf arsenic content (FL-As), Cargo grain arsenic content (CG-As), and the CG-As/FL-As ratio, in the reference (RP) and validation (VP) populations.

**Table 1:**
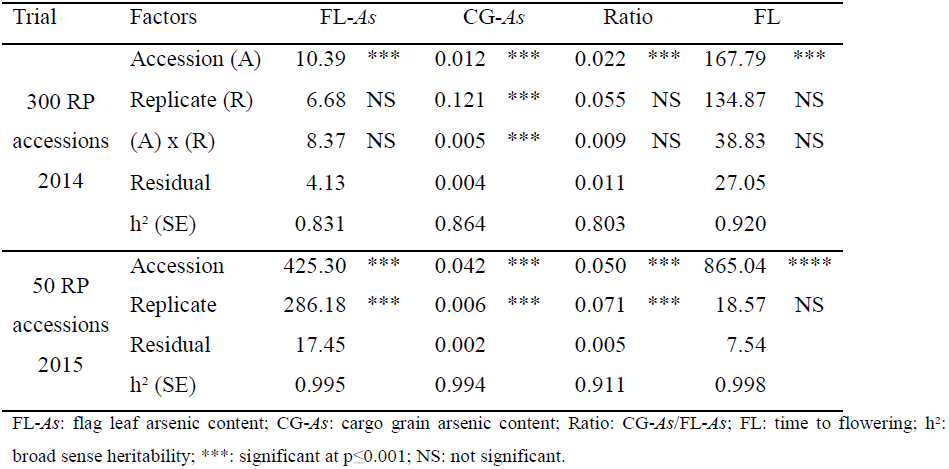
Variance components of three phenotypic traits in the reference population (RP) evaluated in 2014 and in 50 selected accessions of RP evaluated in 2015

| Trial | Factors | FL-As |  | CG-As |  | Ratio |  | FL |  |
| --- | --- | --- | --- | --- | --- | --- | --- | --- | --- |
| 300 RP accessions 2014 | Accession (A) | 10.39 | *** | 0.012 | *** | 0.022 | *** | 167.79 | *** |
|  | Replicate (R) | 6.68 | NS | 0.121 | *** | 0.055 | NS | 134.87 | NS |
|  | (A) x (R) | 8.37 | NS | 0.005 | *** | 0.009 | NS | 38.83 | NS |
|  | Residual | 4.13 |  | 0.004 |  | 0.011 |  | 27.05 |  |
|  | h<sup>2</sup> (SE) | 0.831 |  | 0.864 |  | 0.803 |  | 0.920 |  |
| 50 RP accessions 2015 | Accession | 425.30 | *** | 0.042 | *** | 0.050 | *** | 865.04 | **** |
|  | Replicate | 286.18 | *** | 0.006 | *** | 0.071 | *** | 18.57 | NS |
|  | Residual | 17.45 |  | 0.002 |  | 0.005 |  | 7.54 |  |
|  | h<sup>2</sup> (SE) | 0.995 |  | 0.994 |  | 0.911 |  | 0.998 |  |

FL-As: flag leaf arsenic content; CG-As: cargo grain arsenic content; Ratio: CG-As/FL-As; FL: time to flowering; h<sup>2</sup>: broad sense heritability; \*\*\*: significant at p≤0.001; NS: not significant.

In 2014, variation in FL-*As* among the 300 accessions of RP ranged from 1.34 to 15.61 and averaged 5.88 mg kg^−1^ of dry weight. Variation in CG-*As* ranged from 0.147 to 0.656 mg kg^−1^ and averaged 0.335. The determination coefficient between FL-*As* and CG-*As* was rather low but highly significant (R^2^ = 0.20, p < 0.0001). This rather loose relationship between FL-*As* and CG-*As* corroborates the significant accession effect observed for the CG/FL-*As* ratio.

In 2015, the range of variation in FL-*As* among the 50 accessions of RP with contrasted arsenic contents in 2014 was much larger (from 3.69 to 34.69; average of 16.83 mg kg^−1^), while the range of variation in CG-*As* was slightly narrower (0.169 to 0.493; average of 0.338 mg kg^−1^). However, these differences in the range of variation did not change either the relative ranking of the 50 accessions observed in 2014, or the determining effect of FL-*As* on CG-*As*. Indeed, the Spearman coefficient of rank correlation between performances of the 50 RP accessions in 2014 and 2015 was r = 0.72 (p < 0.0001) for FL-*As*, r = 0.68 (p < 0.0001) for CG-*As*, and r = 0.59 (p < 0.0001) for CG/FL-*As*. Likewise, the determination coefficient between FL-*As* and CG-*As* of the 50 accessions in 2015 was higher (R^2^ = 0.56, p < 0.0001) than the one observed in 2014 for the 300 accessions of RP.

#### Variation in arsenic content in the validation population

Variation in FL-*As* among the 95 accessions in the VP ranged from 3.24 to 37.76 and averaged 14.61 mg kg^−1^. Variation in CG-*As* ranged from 0.208 to 0.729 mg kg^−1^ and averaged 0.341. The determination coefficient between the FL-*As* and CG-*As* was low but highly significant (R^2^ = 0.20, p < 0.0001). The CG-*As*/FL-*As* ratio varied between 0.179 and 0.636 and averaged 0.336 (Figure 1).

### Genetic diversity and structure of the reference and validation populations

Analysis of genetic diversity was performed for 228 RP accessions and 95 VP accessions for whom sufficient GBS data were available for association analysis and genomic prediction.

The 22,370 SNP markers of the working dataset were unevenly distributed along the chromosomes (S1 Figure; S3 Table). Average marker density was one SNP every 17.1 kb. However, it ranged from one SNP every 10.7 kb in chromosome 11 to 26.7 kb in chromosome 9. The number of pairs of loci with a distance greater than 250 kb, 500 kb and 1 Mb was 175, 27 and one, respectively.

The decay of LD over physical distance in the two populations is presented in Figure 2. For between-marker distances of 0 to 25 kb, the average r^2^ was 0.67 and 0.73 in RP and VP, respectively. In the RP, the r^2^ value dropped to half its initial level at around 450 kb, reached 0.2 at 1.25 Mb, and below 0.1 at 2.10 Mb. In the VP, r^2^ reached the 0.2 threshold only at pairwise distances of around 1.70 Mb, and the 0.1 threshold at distances above 3 Mb. No major difference in LD decay was observed between chromosomes. Given these extents of average LDs, one would not expect marker density and distribution along the chromosome to be a major limiting factor for the detection of significant associations and for the predictive ability of genomic prediction.

**Figure 2:**
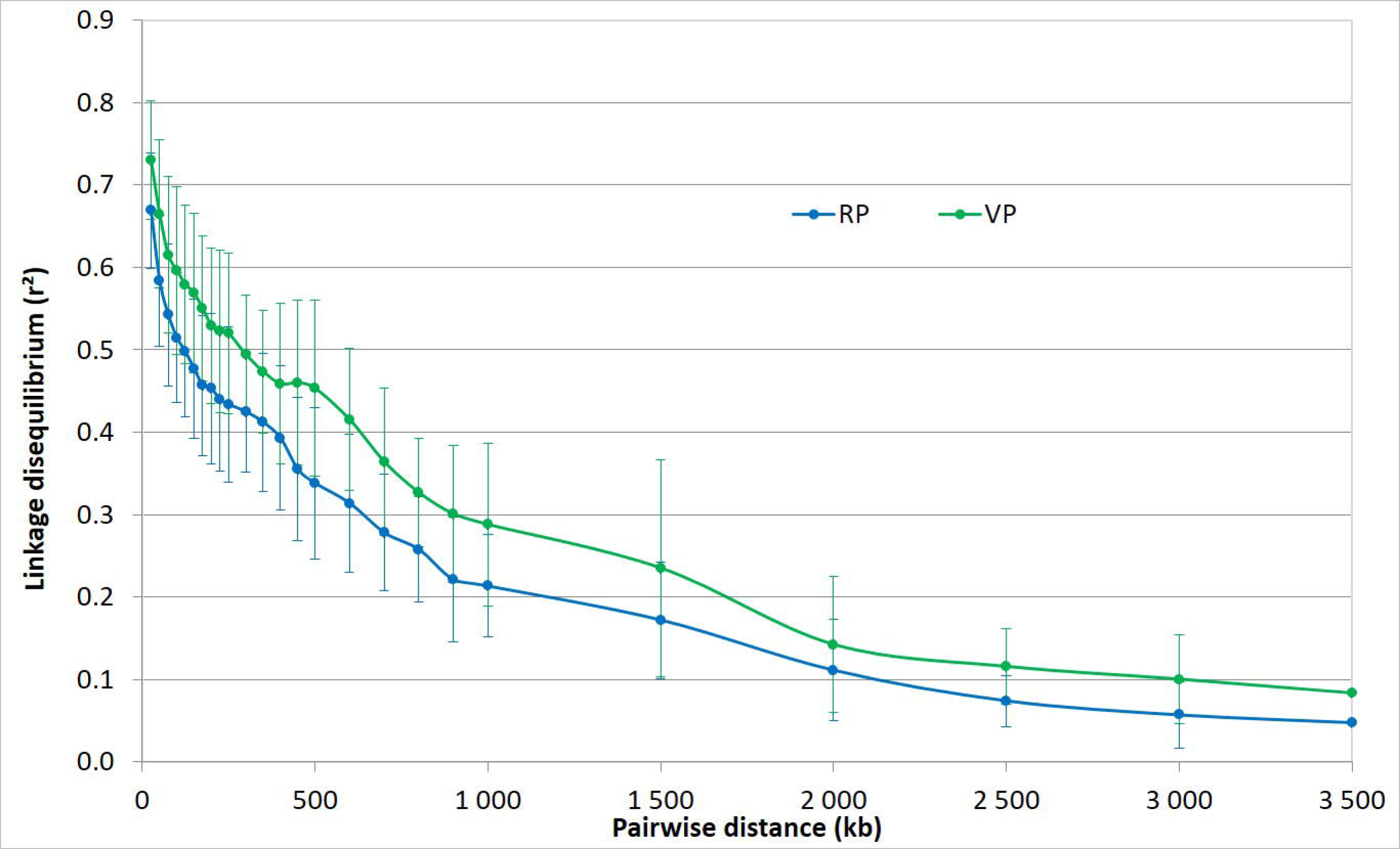
Patterns of decay in linkage disequilibrium in the reference population (green) and in the validation population (blue). The curve represents the average r^2^ among the 12 chromosomes; the bars represent the associated standard deviation.

The two populations showed similar MAF patterns for the 22,370 common SNP loci. RP and VP had the same minor allele in 95.4% of the common loci. In both populations, the MAF distribution was slightly skewed toward low frequencies, the average MAF was close to 22.2%, and the proportion of loci with MAF < 10% was close to 75%. Likewise, the Spearman correlation between the MAF of the 21,343 loci with identical minor alleles in the two populations was r = 0.85 (p <0.01).

Dissymmetry-based clustering of RP accessions led to two major clusters corresponding to the temperate *japonica* (65% of accessions) and tropical *japonica* (35% of accessions) sub-groups (Figure 3). The majority of the temperate *japonica* accessions are of European origin. The majority of the tropical *japonica* accessions originate from the American continent. The inclusion of the VP lines in the analysis did not modify the clustering into two groups. Indeed, 69% of VP lines clustered with the temperate *japonica* group and the remaining 31% with the tropical *japonica* group (Figure 3; S1 Table).

**Figure 3:**
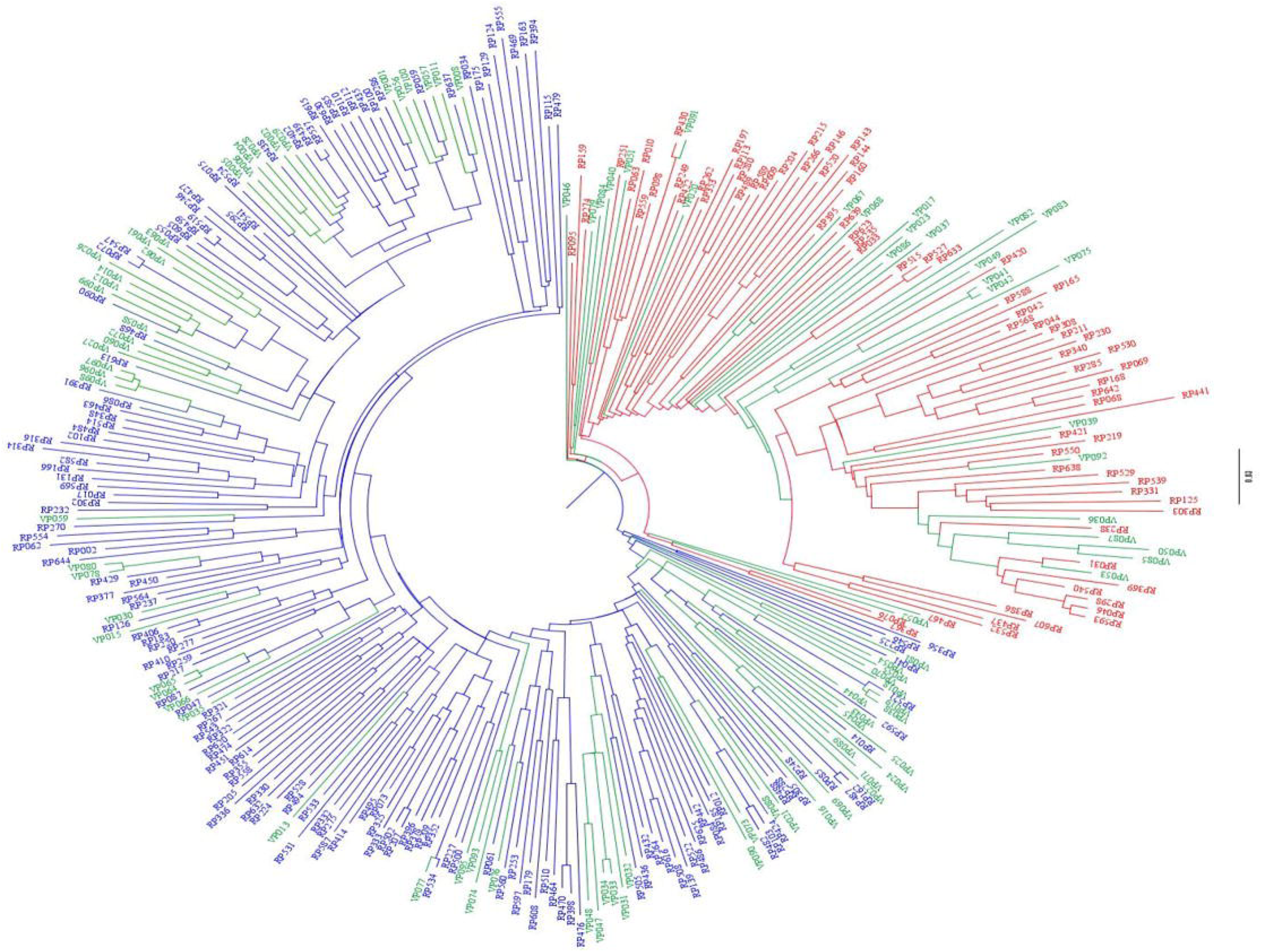
Unweighted neighbor-joining tree based on simple matching distances constructed from the genotype of 228 accessions of the reference population (RP) and 95 advanced lines of the validation population (VP), using 3,620 SNP markers. Green: VP; Red and blue: RP accessions belonging to tropical *japonica* and temperate *japonica*, respectively.

### Relationship between genotypic and phenotypic diversity

Highly significant differences in *As* content were observed between the temperate *japonica* and the tropical *japonica* accessions of RP evaluated in 2014. The former subgroup had the highest arsenic contents (S4 Table; S2 Figure). Data from the 50 RP accessions evaluated in 2015 confirmed this trend. Interestingly, similar to the RP, significant differences in FL-*As* and CG-*As* were also observed between the temperate *japonica* and the tropical *japonica* components of VP, the former subgroup having the highest contents. This superposition of genotypic and phenotypic diversity may negatively influence QTL detection.

### Association analyses

#### Association analysis in the reference population

Results of association analysis of the three traits in the RP are presented in Figure 4 and S5 Table. The number of significant associations (p-value < 1e-05) was 41 for FL-*As*, 23 for CG-*As* and 82 for Ratio. These associations represented 6, 13 and 19 independent loci, i.e. a cluster of SNPs with a distance of less than 1.25 Mb between two consecutive significant SNPs, corresponding to the average LD of r^2^ < 0.2. These loci were composed of 1-35 SNPs, not always adjacent, with p-values ranging between 1e-05 and 1e-07. None of the significant SNPs or independent loci for one trait were found to be significant for another trait. The MAF of the significant SNPs ranged from 2.5% to 49.4% and averaged 36.1% for FL-*As*, 11.7% for CG-*As* and 27.5% for Ratio. The contribution of individual significant SNPs to the total variance of the trait considered (marker R2) was low and did not exceed 12%. Among the 41 SNPs significantly associated with Pl-*As*, 11 corresponding to three independent loci had marker R2 > 10%. The highest marker R2 observed among the 23 SNPs significantly associated with CG-*As*, was 8%. Among the 82 SNPs significantly associated with Ratio, nine corresponding to six independent loci had marker R2 > 10%.

**Figure 4:**
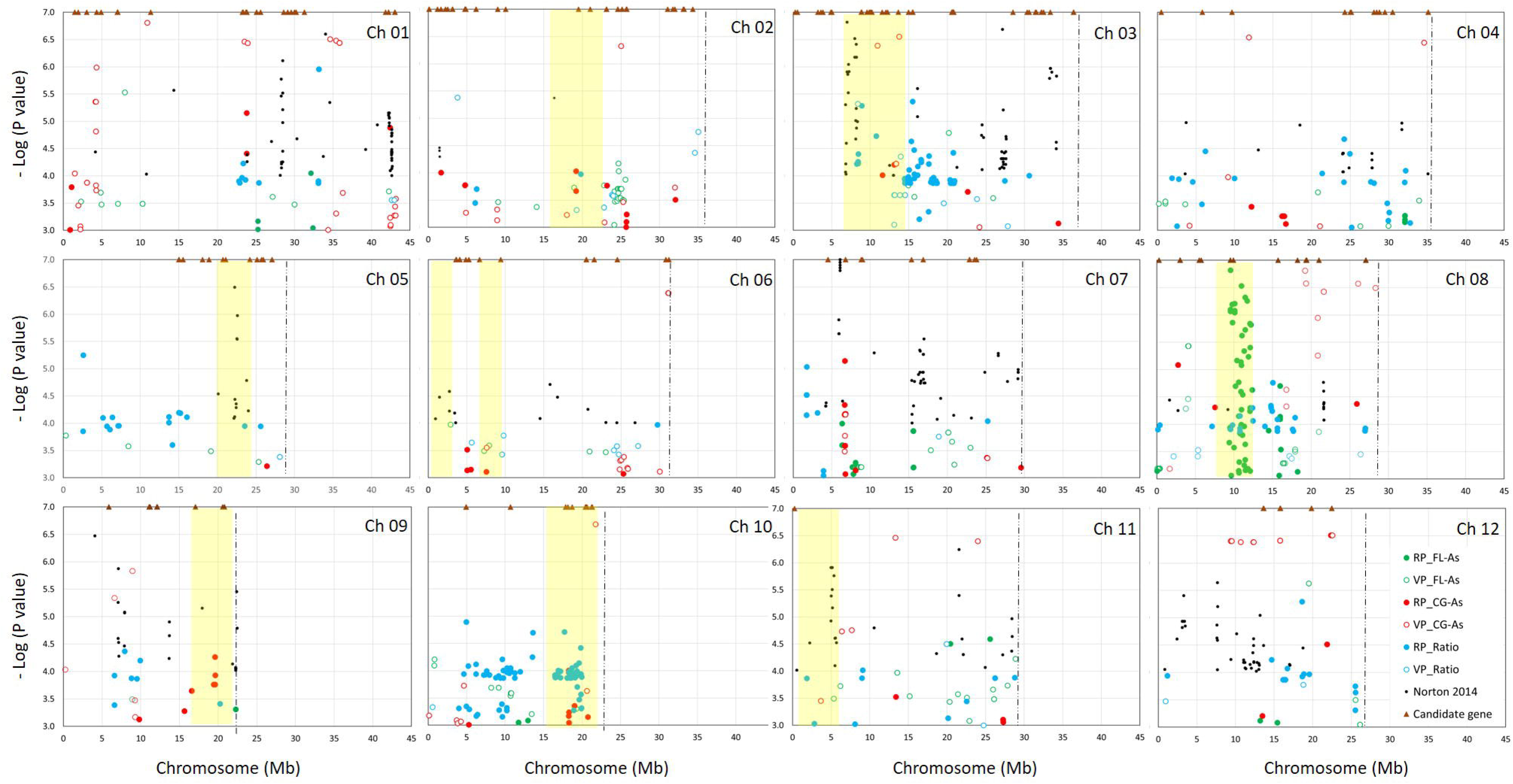
Results of association analyses in the reference population (RP) and the validation population (VP) in the present study, and comparison with data from the literature. For the present study, data points represent SNPs significantly associated with arsenic concentration in the flag leaf (FL-As) in the cargo grain (CG-As), and the CG-As/FL-As ratio in RP and VP. Data from the literature include significant SNPs mapped by GWAS [30], QTLs for grain arsenic concentration [25, 29] and candidate genes [21, 39].

#### Association analysis in the validation population

Results of association analysis for the three traits in the VP are presented in Figure 4 and S5 Table. The number of significant associations was 15 for FL-*As*, 75 for CG-*As* and 8 for Ratio. These associations represented 8, 30 and 5 independent loci. These loci were composed of 1-22 not always adjacent SNPs, with p-values ranging between 1e-05 and 1e-09. Similar to RP, significant SNP loci for the three traits did not colocalize. The MAF of the significant SNP ranged from 2.6% to 46.8% and averaged 28.0% for FL-*As*, 9.0% for CG-*As* and 9.1% for Ratio. The significant SNPs contributed much more, on average, to trait total variance than the ones observed in the RP. The mean marker R2 was 18% for SNPs associated with FL-*As*, 24% for SNPs associated with CG-*As* and 16% for SNPs associated with Ratio.

#### Congruence between the results of GWAS in RP and in VP

Among the 146 SNPs significantly associated with one of the three traits in the RP, only eight were also significant in the VP. These SNPs corresponded to one independent locus associated with CG-*As*. The application of a margin of tolerance of 1.7 Mb between a significant locus in RP and its counterpart in VP (corresponding to the average distances for LD of 0.2 in the VP) only slightly increased the number of colocalizations: four additional colocalizations for CG-*As* and one for Ratio. On the other hand, the number of such colocalizations increased markedly (9, 20 and 12 for FL-*As*, CG-*As* and Ratio, respectively) when the threshold of significance of association in the two populations was lowered to a p-value < 1e-04 (Figure 4). The latter features represented 69%, 40% and 52% of the independent significant loci detected in RP for FL-*As*, CG-*As* and Ratio, respectively.

Genomic localization and co-localization with QTLs and gene reported in the literature

Out of a total of 146 SNPs significantly associated with one of the three *As* related traits in the RP, 41% were located in intergenic regions, 14% in introns, 27% in exons with synonymous coding effects, 10% in exons with non-synonymous coding effects, 6% in UTR-3 regions and 2% in stop-gained sites (S6 Table). The proportions were similar for the 96 significant loci in the VP and for those observed among all the 22,370 SNPs used for GWAS. Genes underlying the significant loci included ATP binding cassette involved in arsenic detoxification (e.g. Os04g0620000), transporters (e.g. phosphate, ammonium, peptide, efflux transporters MATE) abiotic stress responsive genes (e.g. several F-box and DUF domain containing proteins, cytochrome P450) and transcription factors (e.g. MBY, zinc finger family protein, ERF).

A genome survey within an interval of 400 kb (200 kb downstream and 200 kb upstream) surrounding each significant SNP in the RP and in the VP led to the identification of at least one gene with the product involved in plant response to abiotic stresses or reported in the literature as responsive to *As* stress (Figure 4 and S6 Table). The latter included OsLsi1, OsHAC1, OsHAC6, OsACR2-1 and representative of glutathione S-transferases, Cytochrome P450s, heat shock proteins, metal-binding proteins, phosphate acquisition proteins, transporter proteins and transcription factors. Likewise, a survey of the surrounding interval of 400 kb of the significant SNPs for QTL reported in the literature to be associated with *As* resulted in a large number of colocalizations (Figure 4 and S6 Table)

### Genomic prediction

#### Cross validation experiment in the reference population

Application of seven cross validation experiments (corresponding to seven prediction methods) to each of the three phenotypic traits led to average prediction accuracies of 0.484 for FL-*As*, 0.574 for CG-*As* and 0.414 for Ratio (Table 2). Differences in predictive ability between the three traits were highly significant (P < 0.0001). Among the seven prediction methods, RKHS showed the highest average predictive ability (0.475) and BayesB and BayesC the lowest (0.435). However, a marked interaction was observed between prediction methods and traits (Table 2).

**Table 2:** Predictive ability (r) of seven methods of genomic prediction for three rice arsenic content traits in the reference population, based on cross validation experiments.

| Prediction<br>method | Phenotypic traits |  |  |  |  |  | Average<br>r |
| --- | --- | --- | --- | --- | --- | --- | --- |
|  | FL-As |  | CG-As |  | Ratio |  |  |
|  | r | sd | r | sd | r | sd |  |
| GBLUP | 0.449 | 0.155 | 0.535 | 0.166 | 0.356 | 0.159 | **0.446** |
| BayesA | 0.452 | 0.154 | 0.537 | 0.171 | 0.366 | 0.158 | **0.452** |
| BayesB | 0.425 | 0.450 | 0.533 | 0.164 | 0.348 | 0.165 | **0.436** |
| BayesC | 0.442 | 0.160 | 0.519 | 0.163 | 0.344 | 0.162 | **0.435** |
| BL | 0.455 | 0.153 | 0.526 | 0.166 | 0.353 | 0.160 | **0.445** |
| BRR | 0.455 | 0.153 | 0.536 | 0.167 | 0.356 | 0.162 | **0.449** |
| RKHS | 0.408 | 0.941 | 0.549 | 0.086 | 0.468 | 0.125 | **0.475** |
| Average | **0.441** | **0.309** | **0.533** | **0.154** | **0.370** | **0.156** | **0.448** |

FL-As: flag leaf arsenic content; CG-As: cargo grain arsenic content; Ratio: CG-As/FL-As. r: average predictive ability; sd: standard deviation.

In order to evaluate the effect of exclusion of highly redundant SNP (r^2^ = 1), the cross validation experiment was also implemented with the full set of SNPs available (22,370), under GBLUP. Results showed negligible effects on predictive ability: r = 0.449 versus 0.450 with the incidence matrix of 16,902 for FL-*As*, r = 0.535 versus 0.536 for CG-*As*, and r = 0.356 versus 0.357 for Ratio.

#### Genomic prediction across populations

Under the S1 scenario, using all the 228 accessions of the RP as the training set, the predictive ability of genomic estimate of breeding value (GEBV) of the 95 lines of VP was on average 0.426 for FL-*As*, 0.476 for CG-*As* and 0.234 for Ratio (Figure 5 and S7 Table). The three prediction methods implemented provided similar levels of average predictive ability. However, there was some interaction between prediction methods and phenotypic traits. Like for the cross validation experiments, the addition of the redundant SNPs in the incidence matrix did not noticeably modify the predictive ability (Figure 5).

**Figure 5:**
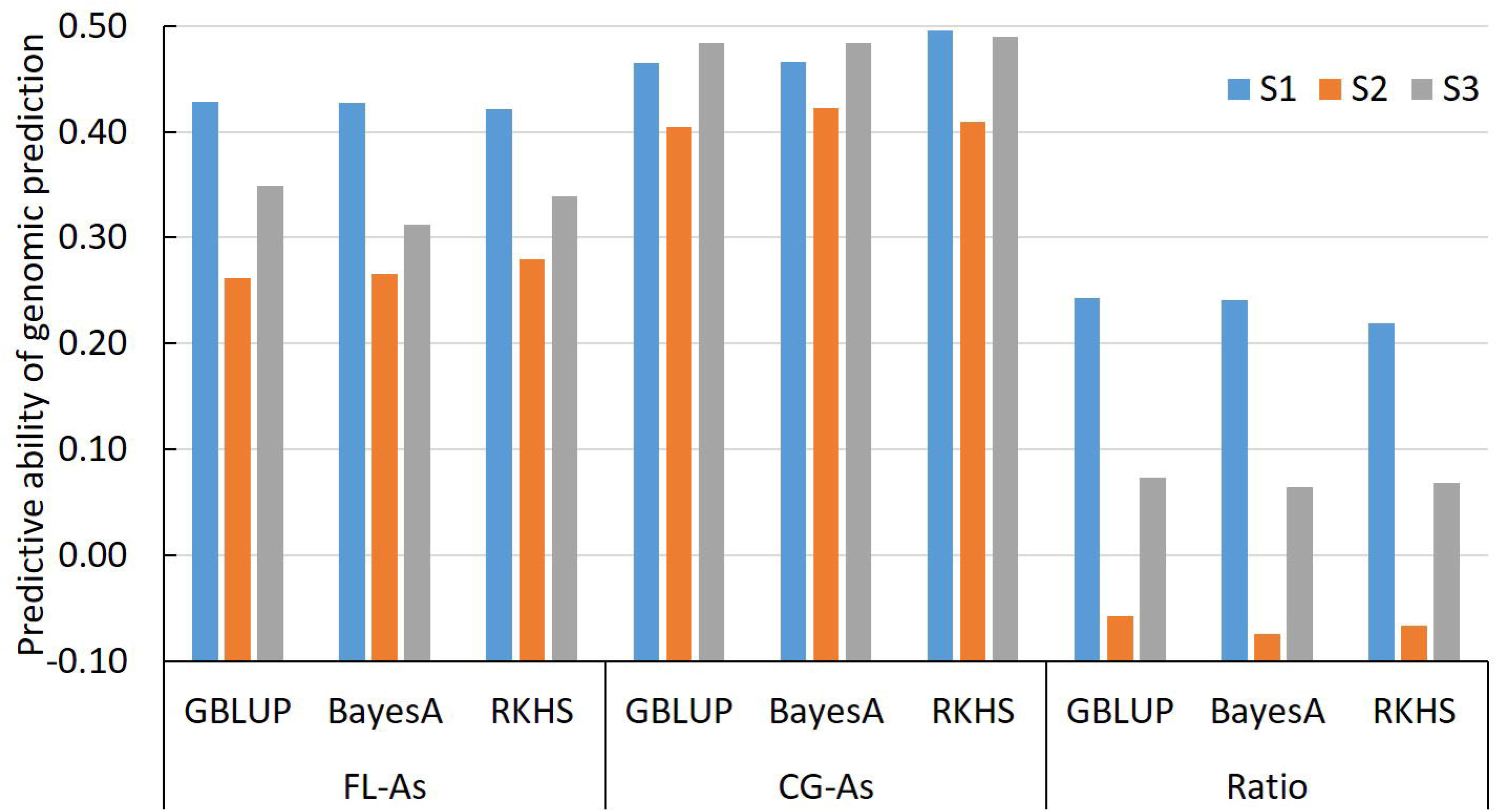
Predictive ability of genomic prediction of the arsenic concentration in the flag leaf (FLAs) in the cargo grain (CG-As), and for the CG-As/FL-As ratio of the validation population obtained with three statistical methods, BayesB, GBLUP and RKHS, under three scenarios of composition of the training set.

The predictive ability of GEBV were much lower under S2, with averages of 0.266, 0.411 and −0.016 for FL-*As*, CG-*As* and Ratio respectively (Figure 5). Under S3, the average predictive ability was slightly higher than under S1 for CG-*As* (0.491), and much lower than under S1 for FL-*As* (0.341) and for Ratio (0.073).

## Discussion

The aim of this work was to explore (i) the phenotypic diversity of the rice *japonica* subspecies, adapted for cultivation in Mediterranean Europe, to restrict *As* accumulation in the grains, and (ii) the potential of the two major options for marker-assisted selection for the improvement of the trait, i.e. QTL-based selection and genomic estimate of breeding value (GEBV)-based selection.

Phenotypic diversity for *As* accumulation was evaluated in field experiments with uncontrolled intensity of exposure to *As*. However, we observed a rather stable soil *As* concentration of about 10 mg kg^−1^ across the crop cycles, and in the three consecutive years of field experiments. This concentration corresponded to the class of rather high *As* contents reported for paddy fields in countries including Bangladesh [37], China [4] and the USA [38]. The range of variation of CG-*As* (0.15 to 0.66 mg kg^−1^) among the accessions of RP was similar to the range observed by [34] in a panel of some 400 accessions representative of the diversity of all the *O. sativa* species (http://www.ricediversity.org/), evaluated in a multilocal trial in Bangladesh, China and the USA. The rather loose relationship between FL-*As* and CG-*As* we observed suggests there are differences between accessions in the ability to limit *As* transfer from the leaves to the grains. To our knowledge, the existence of such genetic diversity for the CG-*As*/FL-*As* ratio has not yet been reported in the literature. The number of rice accessions studied by [37] and [38], who investigated the relationship between rice shoot and grain *As*, was probably too low to reveal the genetic diversity we observed for CG-*As*/FL-*As* ratio. The rather high correlation between the performances of the 50 RP accessions evaluated twice, in two consecutive years, is evidence for the robustness of our findings concerning the extent of genetic diversity for FL-*As* and CG-*As* and on the relationship between the two traits. Interestingly, the extent of FL-*As* and CG-*As* in the VP was as large as that observed in the RP, despite its much smaller size, with only 95 accessions.

In order to explore the potential of marker-QTL association-based breeding for aptitude to restrict *As* accumulation in the grains, we performed association analysis in the RP to detect QTLs. A large number of QTLs was detected for each of the three traits considered. Some of these QTLs colocalized with already reported QTLs [26–29], candidate genes [21, 39], or cloned genes [20, 36]. However, only a few of the QTLs that we detected for the three traits colocalized with each other, some QTLs stretched over several Mb due to the large extent of LD, and none explained more than 10% of total phenotypic variance.

Several factors affect the success of GWAS in precisely mapping QTLs. These include the architecture and the heritability of the target trait, the size and the structure of the population, the number of loci affecting the traits that segregate in the population and their relationship with population structure, the statistical method, and the stringency of the threshold to declare association significance [40]. Apart from the choice of the statistical method and the significance threshold, the experimenter has often limited control over such factors. The exact MLM method we used is known to successfully correct for population structure and family relatedness [41]. Regarding the threshold of significance, several methods have been proposed to overcome the problem of multiple testing. These include monitoring of the number of false positives [42], permutation and boost-trap testing [43], comparing the results of 2-3 different GWAS methods [44], and sub-sampling [45]. However, the only evidence that a significant association detected in a GWAS is “real”, is its validation in an independent population [46], and such a replication requires a sufficiently large validation population to ensure detection power, and with similar features to the initial study of the above-mentioned factors that affect QTL detection [47].

GWAS with our VP detected a similarly large number of SNP and independent loci as with the RP, despite its smaller size (95 entries = 42% of RP size). However, only a few of the QTLs detected in the VP colocalized with the QTLs detected in the RP, despite considerable loosening of the interval surrounding each QTL, or lowering the significance threshold from 1e-05 to 1e-04. Yet, VP had similar features to RP for some of the factors that affect the GWAS results, such as population structure (composed of temperate and tropical *japonica*), the relationship between population structure and variability of the target trait (the temperate *japonica* having the highest *As* contents) and MAF distribution. Likewise, almost all the 95 advanced lines of VP were derived from crosses between members of RP.

Given the above-mentioned superposition of the distributions of the phenotypic variability and the structuring of RP and VP into temperate and tropical *japonica*, our GWAS results might have been subject to an abnormal rate of false negatives due to a confounding phenomenon [48]. To evaluate this risk, we performed separate association analyses with the 153 temperate and the 75 tropical *japonica* accessions of RP. These analyses detected, at best, 50% of the QTLs detected with the entire RP, without markedly increasing the P-value for each association (data not shown). The expected positive effects of diverting the confounding phenomenon proved to be smaller than the reduced detection power due to the reduced size of the population.

The conclusions we draw from these results are that (i) a diversity panel with a large extent of LD has limited genetic resolution power, (ii) it is unlikely that a single GWAS makes it possible to establish robust and precise genotype–phenotype associations, especially for complex traits, and (iii) implementation of an independent replication experiment is a complex process with uncertain results.

To explore the potential of genomic prediction options for breeding for the ability to restrict *As* accumulation in grains, we tested a large set of prediction methods using the cross-validation approach in the RP, and then performed prediction across populations with a smaller set of methods. The level of predictive ability for FL-*As* and CG-*As* in the cross validation experiments was similar to the levels reported in the literature for traits of equivalent heritability in rice [49] and other major crops [50, 51]. Predictions were less accurate for the Ratio trait, which, by design, accumulated the experimental noises associated with the evaluation of FL-*As* and CG-*As*. The cross validation experiments also confirmed the limited differences in predictive ability between prediction methods reported in rice [49, 52] and in other crops [53, 54]. The exclusion of the most redundant SNP markers, based on LD information, had a limited effect on predictive ability, confirming the fact that accounting for LD in the population matters more than the absolute marker density [55].

Across population genomic prediction with models trained with RP data led to slightly lower predictive ability than the predictive ability observed in the cross-validation experiments. Similar decreases in predictive ability have been reported in rice [49], sugar beet [56], barley [51] and strawberry [57], and were attributed to differences in LD and allele frequencies between the training and the validation sets. Differences in the extent and pattern of LD between the training sets represented by diversity panels and the validation sets composed of advanced lines are inevitable [58]. On the other hand, in our case, no significant differences in predictive ability were found between the GBLUP model that captures marker-based relationship between RP and VP, and RKHS and BayesB that captures LD between markers and QTLs. An attempt to reduce the discrepancy in allele frequency between RP and VP by discarding SNP loci with highly divergent MAF did not markedly change predictive ability (data not shown). Neither could conclusive improvement in predictive ability be achieved by optimizing the composition of the training set using the CD-mean approach [59]. These findings suggest that further research aimed at improving the predictive ability of across population genomic predictions should explore the effects of the size of the training set (use a larger training set) and of the balance between marker density and the regularity of their distribution along the genome. Indeed, in the present work, marker density (one SNP every 17.1 kb) was rather high, given the extent of LD, but their distribution was not optimized given the GBS genotyping technology.

The critical importance of reducing the presence of *As* in the rice grains in a large proportion of rice growing areas has recently resulted in steady efforts to understand the molecular mechanisms involved in plant response to overexposure to *As* [10, 22] and the genetic control of these mechanisms [15, 16, 19]. Although a few genes, reported as being “crucial”, have been cloned [36], transcriptome analyses [21, 39] and GWAS results [30] suggest that *As* tolerance is a complex trait involving a large number of loci with limited individual effect on the trait.

The number of candidate loci makes marker-assisted pyramiding of the favourable alleles unpractical. Moreover, uncertainty concerning the exact genomic position of some of the loci makes the outcome of marker-assisted pyramiding unpredictable. Indeed, as discussed above, GWAS results raise robustness issues, and this also seems to be the case for transcriptome analyses [60].

The GEBV we obtained for flag leaf and cargo grain *As* contents were reasonably accurate in both intra-population (cross validation in the RP) and across-population (RP/VP) prediction experiments. Translation of those prediction accuracies into average phenotypic performances of VP lines selected based on their GEBV by model trained with the RP is even more encouraging. Indeed, the average FL-*As* and CG-*As* of the best 10 VP lines selected on the basis of phenotypic data were 41% and 65% of the average FL-*As* and CG-*As* of all 95 lines of VP. The average FL-*As* and CG-*As* of the best 10 VP lines selected on the base of GEBV were 55% and 85% of the average FL-*As* and CG-*As* the whole 95 lines of VP (S8 Table). In other words, for a selection rate of 10%, the difference in genetic gain between phenotypic selection and GEBV based selection was only 10% for FL-*As* and 5% for CG-*As*. Given these rather small differences in genetic gains, the choice between phenotypic and GEBV based selection will depend mainly on the comparative costs of genotyping and phenotyping for *As* content. If the costs are similar, the best choice would be GEBV-based selection because genotypic data are a multi-purpose asset that can also be used for genomic prediction of other traits than *As* content. The possibility of changing the genotyping method to obtain a smaller but more evenly distributed number of markers should also be considered in the decision making process. Indeed, simulation works [61] and experimental data [42] have shown that, if markers are chosen based on LD distribution along the chromosomes, the number of markers can be reduced drastically without affecting predictive ability.

To conclude, considering the limitations of QTL-based marker-assisted selection for *As* and the level of predictive ability of GEBV, genomic prediction proves to be the most promising option for breeding for the ability to restrict *As* accumulation in the rice grain. In a previous study [49], we showed that a rice diversity panel could provide accurate genomic predictions for complex traits in the progenies of biparental crosses involving members of the panel. In addition, associated with the rapid generation advancement technique, genomic selection can accelerate the genetic gain of the pedigree breeding scheme, the most common breeding scheme in rice. GS for *As* content can be incorporated in such a breeding program. The main additional cost would be the phenotyping of the diversity/reference panel for *As* content.

## Methods

### Plant material

The initial plant material comprised a diversity panel of 300 accessions and set of 100 advanced inbred lines (F5–F7), all belonging to the *japonica* subspecies of *O. sativa*, and adapted to cultivation in the irrigated rice ecosystem of temperate Mediterranean Europe. The diversity panel, hereafter referred to as the reference population (RP), was composed of 214 accessions representing the European Rice Core Collection (ERCC), established by merging the working collections of five European public rice breeding programs in France, Greece, Italy, Portugal and Spain [62], and 86 accessions of direct interest for the Camargue-France breeding program (S1 Table). The 95 advanced breeding lines hereafter referred to as the validation population (VP), was composed of elite lines of the rice breeding program run by the *Centre Français du Riz* (CFR) and Cirad, in the Camargue region, France.

### Field trials and phenotyping

Field trials were conducted at the CFR experimental station, Mas d’Adrien (43°42’13.77”N; 4°33’44.71”E; 3 m asl.), under a standard irrigated rice cropping system. The RP was phenotyped in two consecutive years (2014 and 2015), the VP only in 2016. In 2014, all 300 accessions of RP were phenotyped under an augmented randomized complete block design repeated twice, each block being composed of 25 tested accessions and two check varieties (Albaron and Brio). In 2015, 50 accessions of RP, with contrasted *As* content performances, were phenotyped in complete randomized blocks with eight replicates. In both 2014 and 2015 trials, the size of the individual plot was one row of 15 plants. In 2016, each of the 95 advanced lines of VP was represented by five full-sib lines and the size of the individual plot for each full-sib line was one row of 15 plants.

In each field trial, the concentration of total arsenic in the flag leaf (FL-*As*) and in the cargo grain (CG-*As*) was measured and the CG-*As*/FL-*As* ratio calculated. In the 2014 and 2015 trials, three biological samples were prepared for each individual plot to measure FL-*As*. Each biological sample was composed of three flag leaves of three different plants. Each biological sample was oven-dried at 75°C for 120 h, ground, mineralized, and total arsenic concentration was measured using the inductively coupled plasma mass spectrum (ICP-MA; Bruker Aurora ICP Mass Spectrometer). For each biological sample, total arsenic was measured in at least two technical samples and averaged to establish the sample phenotype. Data from the three biological samples were averaged to establish the plot phenotype. A similar procedure was applied to CG-*As* measurement in which the biological samples were composed of three panicles. These panicles were threshed after oven drying, the resulting paddy grains were dehusked, and the cargo grain was ground before undergoing the mineralization procedure.

In 2016, FL-*As* and CG-*As* were measured in one randomly chosen sib-line in each advanced line. Two biological samples were prepared from each chosen sib-line: one biological sample from an individual plant that was also used for DNA extraction and genotyping (see below), and a second sample from the bulk of at least three plants.

In each field trial, the soil total *As* content was measured before sowing and after harvest. Likewise, in each field trial, total *As* content of irrigation water was monitored once a month during the rice cropping cycle.

### Genotypic data

Genotypic data were produced by two distinct genotyping by sequencing (GBS) experiments, for 228 accessions of RP and 95 lines of VP. In both cases, DNA libraries were prepared at the Regional Genotyping Technology Platform (http://www.gptr-lr-genotypage.com) hosted by Cirad, Montpellier France). Genomic DNA was extracted from the leaf tissues of a single plant from each accession using the MATAB method and then diluted to 100 ng/µl. Each DNA sample was digested separately with the restriction enzyme *ApekI*. DNA libraries were then single-end sequenced in a single-flow cell channel (i.e., 96-plex sequencing) using an Illumina HiSeq™2000 (Illumina, Inc.) at the Regional Genotyping Platform (http://get.genotoul.fr/) hosted by INRA, Toulouse, France. The fastq sequences were aligned to the rice reference genome (Os-Nipponbare-Reference-IRGSP-1.0 [63] with Bowtie2 (default parameters). Non-aligning sequences and sequences with multiple positions were discarded. Single nucleotide polymorphism (SNP) calling was performed using the Tassel GBS pipeline v5.2.29. The initial filters applied were the quality score (>20), the count of minor alleles (>1), and the bi-allelic status of SNPs. In the second step, loci with minor allele frequency (MAF) below 2.5% and with more than 20% missing data were discarded. The missing data were imputed using Beagle v4.0. The RP and VP genotyping experiment yielded 39,497 and 67,658 SNP loci, respectively, among which 22,370 were common to the two populations. This working dataset can be downloaded in HapMap format from http://tropgenedb.cirad.fr/tropgene/JSP/interface.jsp?module=RICE study Genotypes, study type ML panel-GBS-data.

### Analysis of phenotypic data

In 2014, RP plot phenotypic data of the 300 accessions were modeled for each trait as:

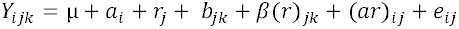

where *Y*_*ijk*_ is the observed phenotype of accession *i* in replicate *j* and bloc *k*, μ is the overall mean, *a*_*i*_ the accession effect, *r*_*j*_ the replicate effect, *b*_*jk*_ the check effect considered as quantitative covariate, *β*(*r*)_*jk*_ the block effect within the replicate, (*ar*)_*ij*_ the interaction between accessions and replicates, and *e*_*ij*_ the residual.

In 2015, RP plot phenotypic data of the 100 advanced lines were modeled for each trait as *Y*_*ij*_ = μ + *a*_*i*_ + *r*_*j*_ + (*ar*)_*ij*_ + *e*_*ij*_ where *Y*_*ij*_ is the observed phenotype of accession *i* in bloc *j*, μ is the overall mean, *a*_*i*_ the accession effect, *r*_*i*_ the replicate effect, (*ar*)_*ij*_ the interaction between accession *i* and replicate *j*, considered as random, and *e*_*ij*_ the residual. For each dataset and each trait, least square means were estimated using the mixed model procedure of Minitab 18.1.0 statistical software (Minitab Inc. 2017).

Broad-sense heritability was calculated for each trait as: 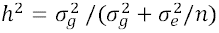, where 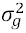 and 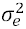 are the estimates of genetic and residual variances, respectively, derived from the expected mean squares of the analysis of variance and *n* is the number of replicates. The computed CG-*As*/FL*-As* ratio were subjected to the angular transformation *2Arcsin square root* before analysis.

### Genotypic characterization of RP and VP

The genetic structure of 228 accessions of RP and 95 advanced lines of VP was analyzed jointly using a distance-based method. First, a matrix of 3,620 SNPs was extracted from the working genotypic dataset of 22,370 SNPs common to RP and VP, by discarding loci that had imputed data and by imposing a minimum distance of 25 kb between two adjacent loci. Then, an unweighted neighbor-joining tree based on dissymmetry matrix was constructed using DarWin v6.

The speed of decay of linkage disequilibrium (LD) in RP and VP was estimated by computing r^2^ between pairs of markers on a chromosome basis using Tassel 5.2 software, and then averaging the results by distance classes using XLSTAT.

### Association analysis

Separate association analyses were performed with phenotypic and genotypic data from 228 accessions of RP and from 95 advanced lines of VP. A single marker regression-based association analysis was performed for each phenotypic trait under a mixed linear model (MLM), in which marker and population structure (Q matrix) effects were considered as fixed and the kinship effect (K matrix) was considered as random. The MLM was run under the exact method option of Tassel 5.2 software, where the additive genetic and residual variance components are re-estimated for each SNP. For each SNP tested, Tassel 5.2 computed a p-value, the log likelihood of the null and alternative models, and the fixed-effect weight of the SNP with its standard error. The threshold to declare the association of a SNP marker with a trait to be significant was set at a probability level of 1e-05. Genes underlying the significant loci were analyzed using the MSU database (http://rice.plantbiology.msu.edu/) search and gene annotation.

### Genomic prediction

#### Construction of the incidence matrix

In order to reduce possible negative effects of redundancy of marker information on the predictive ability of genomic predictions and to reduce computing time, redundant SNPs were discarded as follows. First, using the genotypic dataset of the RP (N = 228 entries and P = 22,370 SNPs), for each SNP, pairwise LD with all other SNPs was calculated. Second, among each group of SNPs in complete LD (r^2^ = 1), the first SNP along the chromosome was maintained and all the others were discarded. This procedure reduced the total number of SNP loci to 16,902. Once the list of these SNPs was established, the incidence matrix of 16,902 SNP was constructed for the VP accordingly.

#### Cross validation experiment in the RP

Seven statistical methods were tested: genomic best linear unbiased prediction (GBLUP), BayesA, BayesB and BayesC, Bayesian lasso and Bayesian ridge regression, and the reproducing kernel Hilbert spaces regressions (RKHS), using the *BGLR* statistical package [64]. The default parameters for prior specification were used and the number of iterations for the Markov chain Monte Carlo (MCMC) algorithm was set to 25,000 with a burn-in period of 5,000.

The cross validation experiments used 171 (3/4) of the 228 accessions of the RP as the training set and the remaining 57 (1/4) accessions as the validation set. Each cross validation experiment was repeated 100 times using 100 independent partitioning of the RP into training set and validation set. For each independent partitioning, the correlation between the predicted and the observed phenotype was calculated so as to obtain 100 correlations for each cross validation experiment. The predictive ability of each cross validation experiment was computed as the mean value of the 100 correlations.

To analyze sources of variation in the predictive ability of genomic predictions, the correlation (*r*) of all prediction experiments was transformed into a Z statistic using the equation: *Z* = 0.5 {*ln*[1 + *r*] − *ln*[1 − *r*]} and analyzed as a dependent variable in an analysis of variance. After estimation of confidence limits and means for Z, these were transformed back to *r* variables.

### Genomic prediction across populations

The predictive ability of genomic prediction across populations was evaluated under three scenarios of composition of the training set. Under the first scenario (S1), all 228 accessions of the RP were used as the training set. Under S2, the training set was composed of the 100 accessions of the RP with the lowest average pairwise Euclidian distances with the 95 lines of the VP. Under S3, 100 accessions of the training set were selected among the 228 accessions of RP, using the CDmean method of optimization of the training set [59]. In this 3rd scenario, a dedicated training set was selected for each phenotypic trait to account for trait heritability. Three statistical methods GBLUP, BayesA and RKHS (that provided the highest predictive ability in the cross-validation experiments) were tested using the *BGLR* statistical package [64]. For each trait, the predictive ability of the prediction experiment was calculated as the correlation between the predicted and the observed phenotypes of the 95 lines.

## Acknowledgements

This work was supported by the CIRAD - UMR AGAP HPC Data Center of the South Green Bioinformatics platform (http://www.southgreen.fr/). We thank the members of CFR who helped conducting the field experiments and the members of the AGAP joint research unit who helped sample leaves and panicles in the field. We thank Brigitte Courtois for her critical review of the manuscript.

## Funding

This work was funded by FranceAgrimer (http://www.franceagrimer.fr/), Grants SIVAL n°2013-1296, SIVAL n° 2014-1382, and SIVAL n° 2015-0761.

## Authors’ contributions

NA: conceived the study, analyzed the data and wrote the manuscript.

JF: produced the genotypic and phenotypic data and wrote the manuscript.

GAS: provided expertise and laboratory facilities for the measurement of arsenic concentration. AL and AB: ran the field experiments and selected the advanced lines composing the validation population.

## Additional information

### Data availability statement

The Phenotypic data analyzed for this study are included in the supplementary table 1

The datasets generated and analysed during the current study are available in the ML panel-GBS-data repository, http://tropgenedb.cirad.fr/tropgene/JSP/interface.jsp?module=RICE

### Competing interests

The authors declare that they have no competing interests

## Supporting information

**S1 Table.** Main characteristics of the 228 accessions of the reference population (RP) and 95 advanced lines of the validation population.

**S2 Table.** Soil and water arsenic contents in the experimental site over the three years of field experiments.

**S3 Table.** Variability of marker density and frequency of minor alleles (MAF) along the 12 chromosomes in the reference and the validation populations.

**S4 Table.** Average arsenic contents of the two subgroups of *O. sativa japonica* present in the reference population (RP) and in the validation population (VP).

**S5 Table.** Results of association analysis of the concentration of arsenic in the flag leaf (FL-As) in the cargo grain (CG-As), and for the CG-As/FL-As ratio, in the reference population (RP) and in the validation population (RV).

**S6 Table.** Colocalization of SNP loci significantly associated with arsenic content traits in the present study with similar loci reported in the literature.

**S7 Table 7.** Predictive ability of genomic estimate of breeding value of the 95 advanced lines of the validation population for arsenic contents, by three genomic prediction models trained with data from 228 accessions of the reference population.

**S8 table 8.** Translation of predictive ability of genomic prediction into genetic gain under different selection intensities.

**S1 Figure.** Distribution of the 22,370 working set SNP markers along the 12 chromosomes in the reference and validation populations.

**S2 Figure.** Distribution of adjusted phenotypic values for arsenic content of the flag leaf (FL-As) and arsenic content of the cargo grain (CG-As), in the reference and validation populations, according to membership of the accessions of temperate japonica and tropical japonica subgroups.

